# Validation of novel *Mycobacterium tuberculosis* isoniazid resistance mutations not detectable by common molecular tests

**DOI:** 10.1101/322750

**Authors:** Justin L. Kandler, Alexandra D. Mercante, Tracy L. Dalton, Matthew N. Ezewudo, Lauren S. Cowan, Scott P. Burns, Beverly Metchock, Global PETTS Investigatorsc, Peter Cegielski, James E. Posey

**Author notes:** Present address: Fulgent Genetics, Atlanta, Georgia, United States (Justin L. Kandler). Present address: Enteric Diseases Laboratory Branch, Division of Foodborne, Waterborne, and Environmental Diseases, National Center for Emerging Zoonotic and Infectious Diseases, Centers for Disease Control and Prevention, Atlanta, Georgia, United States (Alexandra D. Mercante). Please address any correspondence to James E. Posey.

## Abstract

Resistance to the first-line anti-tuberculosis (TB) drug, isoniazid (INH), is widespread, and the mechanism of resistance is unknown in approximately 15% of INH-resistant (INH-R) strains. To improve molecular detection of INH-R TB, we used whole genome sequencing (WGS) to analyze 52 phenotypically INH-R *Mycobacterium tuberculosis* complex (MTBC) clinical isolates that lacked the common *katG* S315T or *inhA* promoter mutations. Approximately 94% (49/52) of strains had mutations at known INH-associated loci that were likely to confer INH resistance. All such mutations would be detectable by sequencing more DNA adjacent to existing target regions. Use of WGS minimized the chances of missing infrequent INH resistance mutations outside commonly targeted hotspots. We used recombineering to generate 12 observed clinical *katG* mutations in the pansusceptible H37Rv reference strain and determined their impact on INH resistance. Our functional genetic experiments have confirmed the role of seven suspected INH resistance mutations and discovered five novel INH resistance mutations. All recombineered *katG* mutations conferred resistance to INH at a minimum inhibitory concentration of ≥0.25 μg/mL and should be added to the list of INH resistance determinants targeted by molecular diagnostic assays. We conclude that WGS is a superior method for detection of INH-R MTBC compared to current targeted molecular testing methods and could provide earlier diagnosis of drug-resistant TB.

## Introduction

Tuberculosis (TB) is the leading cause of death from a single infectious agent (1), yet in 2014, only 3.8% of global development assistance for health funding was for TB (2). The World Health Organization (WHO) estimates that in 2016 alone, 1.674 million people died from TB (1). While ~52.5 million lives were saved between 2000 and 2016 by improving diagnostics and treatments (1), 13 of the 22 highest-burden countries were failing to meet TB prevalence reduction benchmarks in 2014 (3), highlighting the need for enhanced strategies to combat this disease. In the absence of an effective vaccine against pulmonary tuberculosis, suitable antibiotic therapy remains the most important means to stop TB.

Isoniazid (INH) rapidly kills actively growing *Mycobacterium tuberculosis* (*Mtb*) by inhibiting mycolic acid synthesis and thus destroying bacterial cell wall integrity (4) and is a crucial component of drug regimens for both latent TB infection and active TB disease (5, 6). INH remains on the WHO’s lists of essential medicines for both children and adults (7, 8) and can be used for TB therapy even in cases where pregnancy and HIV may be complicating factors (9). However, INH resistance is globally widespread and ranged from ~6%-45% (years 1994-2009), depending on the region (10). Importantly, INH resistance is strongly associated with subsequent treatment failure, relapse, and acquired multidrug resistance (11) and precedes the acquisition of resistance to most other antimycobacterial drugs. The *katG* S315T, *inhA* I194T, and *fabG1-inhA* c-15t mutations arise before all other drug resistance mutations so frequently that they have been deemed “harbinger mutations”, and early detection may be helpful in preventing the evolution and spread of multidrug resistant and extensively drug resistant (MDR and XDR) strains (12). If susceptibility to INH determines the success or failure of TB chemotherapy and the acquisition of resistance to other drugs, future diagnostic tools must strive to optimize sensitivity and specificity in order to limit cases where INH resistance is misdiagnosed.

In *Mtb*, mechanisms of resistance to INH have been studied extensively (reviewed in (13)). Yet, according to a systematic review of publications summarizing genotypic data for 8,796 phenotypically INH-R clinical isolates from 49 countries (14), approximately 15% of INH-R strains could not be explained by known resistance-conferring mutations. By far the most common mutations leading to resistance are the *katG* S315T mutation and the (*mabA*) *fabG1-inhA* c-15t promoter mutation, which confer resistance by preventing activation of the INH prodrug (15, 16) or overexpressing the target of active INH (17), respectively. When combined, these two mutations account for approximately 83% of INH resistance (14). Other rare mutations have been experimentally confirmed as resistance determinants in *Mtb* by functional genetics and include the T275P and W300G mutations in *katG* (16, 18); partial or complete deletion of the *katG* open reading frame (19, 20); the g-7a, a-10c, and g-12c mutations in the *furA-katG* intergenic region (21); deletion of a 134 bp fragment that includes the *furA-katG* promoter (22); an S94A mutation in *inhA* (17); and the L203L silent mutation in *fabG1*, which increases expression of the downstream *inhA* gene (23). Numerous other mutations have been associated with INH resistance in *Mtb* clinical isolates but have yet to be investigated by functional genetics to confirm their role (13). Changes to the promoter region of the *ahpC* gene, encoding an alkyl hydroperoxidase, are frequently found in INH-R clinical isolates of *Mtb* but their role (if any) in directly conferring INH resistance remains unclear (13, 24).

Molecular methods commonly used to detect INH-R TB include line probe assays [LPAs; (25)] and sequencing of known resistance-associated genes by the Sanger method (26) or pyrosequencing (27). The WHO updated its recommendations in 2016 (28) to approve the use of the Hain GenoType MTBDR*plus* version 2 (Hain Lifescience, Nehren, Germany) and Nipro NTM+MDRTB Detection Kit 2 (Nipro, Tokyo, Japan) LPAs. Both tests detect the *katG* S315T and *fabG1-inhA* c-15t/t-8c mutations. Additionally, MTBDR*plus* version 2 detects *fabG1-inhA* a-16g and t-8a mutations, while NTM+MDRTB Detection Kit 2 can detect the *katG* S315N mutation (29, 30). In contrast to LPAs, targeted sequencing allows for customizable mutation detection. For example, the US Centers for Disease Control and Prevention (CDC) offers the <u>M</u>olecular <u>D</u>etection of <u>D</u>rug <u>R</u>esistance (MDDR) service (31), which detects mutations associated with first- and second-line drug resistance. Mutations associated with INH resistance are detected by MDDR by using Sanger and pyrosequencing to target the following: the *katG* S315 codon, the *fabG1-inhA* promoter, the *ahpC* promoter, and the *fabG1* L203 codon.

Despite these molecular tools, phenotypic drug susceptibility testing (DST) is still considered the “gold standard” for determining MTBC drug resistance. Phenotypic DST requires costly infrastructure for culture (e.g. BSL-3 laboratory space etc.) and specialized training and equipment for MTBC-specific susceptibility testing (e.g. Bactec MGIT system). These difficulties, along with slow growth of MTBC cultures and the laborious nature of phenotypic DST assays, have highlighted the utility of molecular tests for detection of MTBC drug resistance. Although rapid molecular tests greatly decrease the turnaround time for resistance prediction compared to phenotypic DST, phenotypic resistance can occur in the absence of known genetic markers (i.e. discordance). Therefore, rapid molecular testing generally complements and does not replace phenotypic DST.

Strains of INH-R MTBC lacking common INH resistance mutations most likely encode resistance elsewhere in the genome and might hold clues to uncover INH resistance mechanisms in up to 15% of MTBC strains (14) whose resistance would not be detected by conventional rapid molecular testing (LPA, Sanger/pyrosequencing). In this work, we used whole genome sequencing (WGS) and phenotypic DST to determine probable causes of INH resistance in clinical isolates of MTBC lacking the common *katG* S315T and *fabG1-inhA* t-8a, t-8c, c-15t, a-16g promoter mutations. We also used functional genetics to assess the impact of several observed *katG* mutations on the INH minimum inhibitory concentration (MIC) of the pansusceptible *Mtb* strain, H37Rv.

## Materials and Methods

### Bacterial strains, plasmids, and oligonucleotides

All clinical isolates that met the criteria for study inclusion (Figure 1) are listed in Table 1 and were evaluated for the presence of the *katG* S315T and *fabG1-inhA* t-8a, t-8c, c-15t, a-16g mutations by Hain Genotype MTBDR*plus* v2 LPA test or Sanger/pyrosequencing prior to this work. To determine the phylogenetic diversity of our study set, we performed single nucleotide polymorphism (SNP)-based analysis of all 52 genomes (Table 1; (32)) using our Unified Variant Pipeline (CDC, unpublished data). Recombineered strains were derived from the H37Rv wild-type parent strain and are listed in **Supplemental Table S1**. Plasmids and oligonucleotides used are listed in **Supplemental Table S2**. pJV128 (33) was provided by Graham F. Hatfull (University of Pittsburgh). All mycobacterial broth cultures were grown as described previously (34). This work was approved by the CDC Institutional Review Board (protocol 5411).

**Figure 1:**
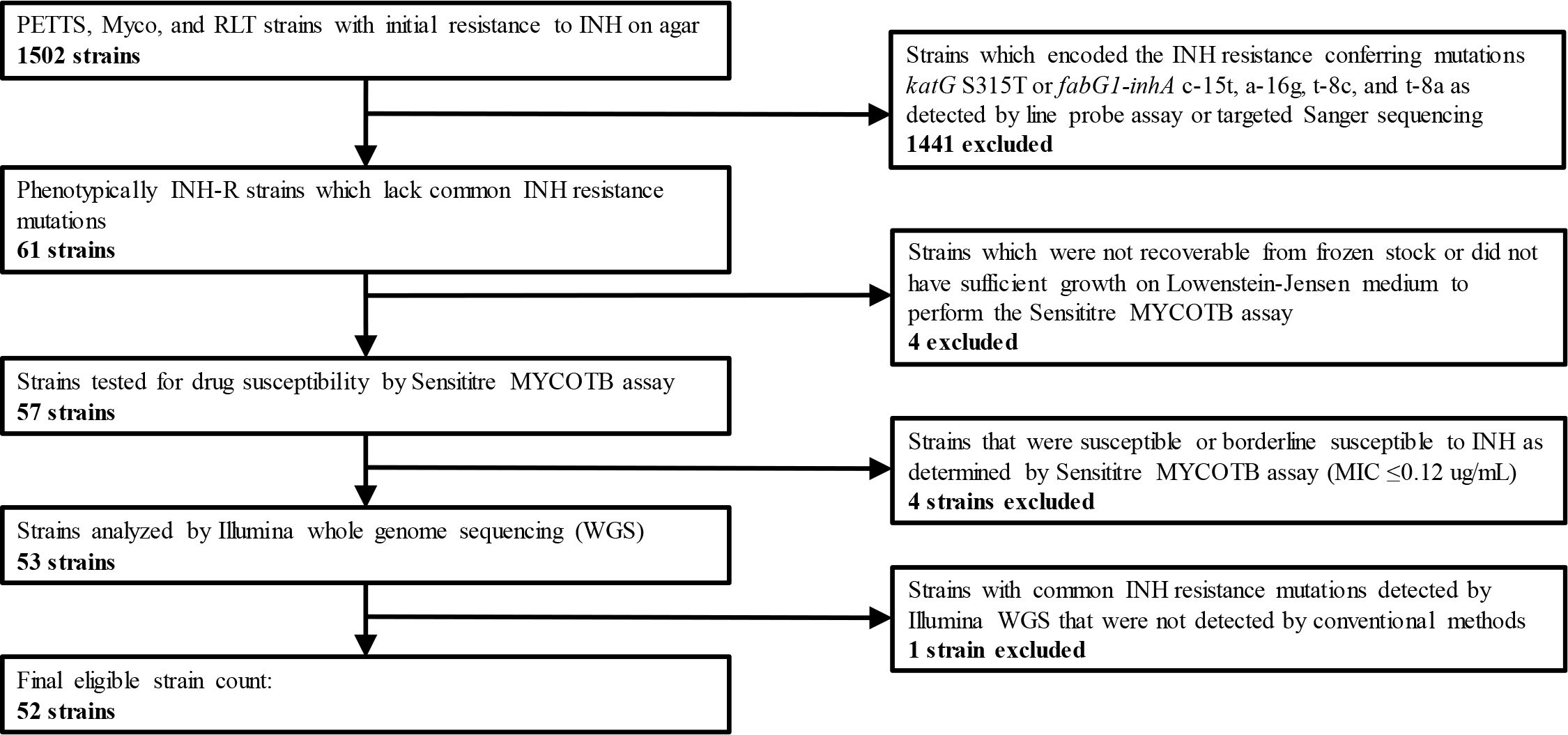
Selection and exclusion criteria for clinical strains included in this study.

**Table 1:**
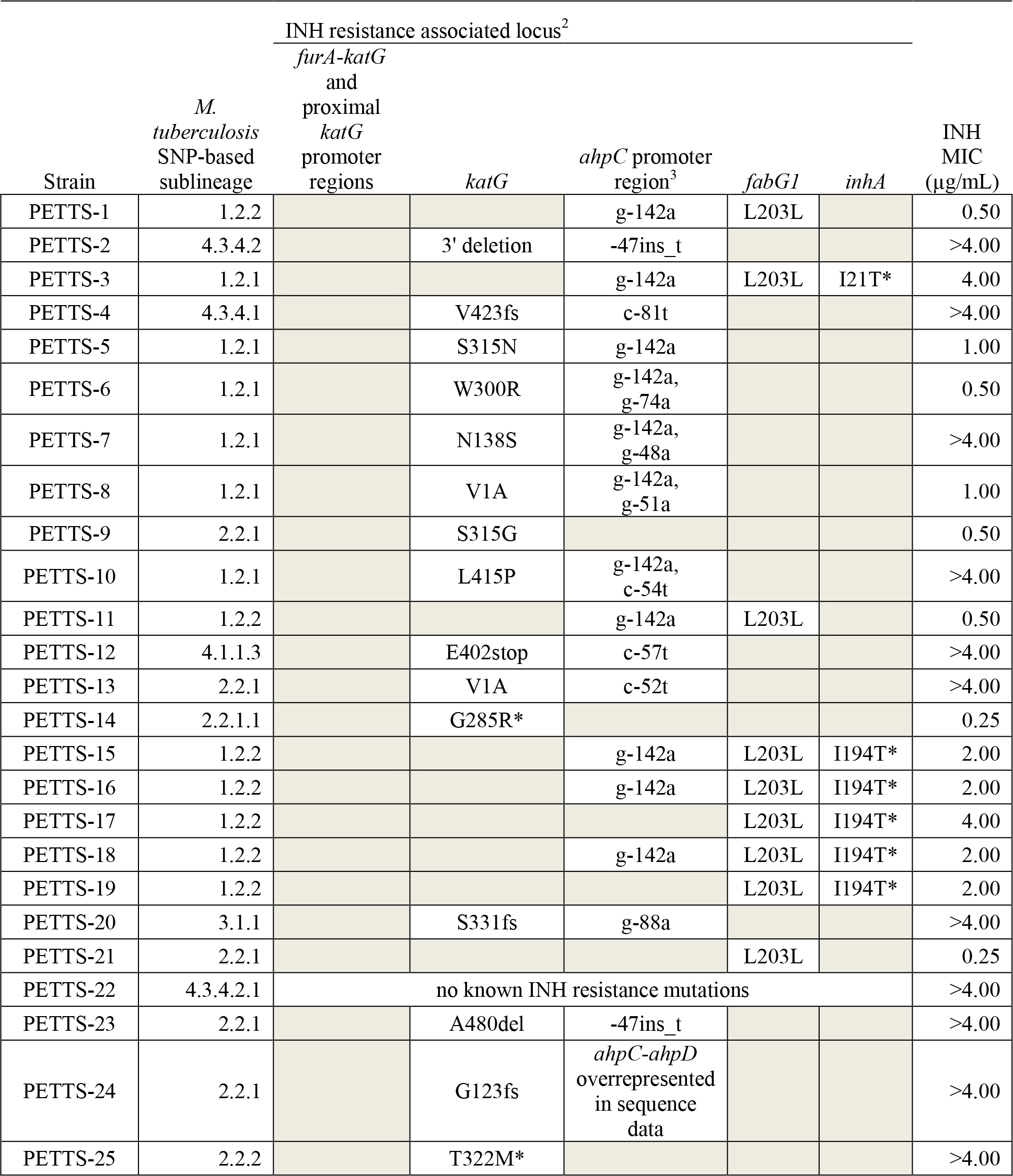
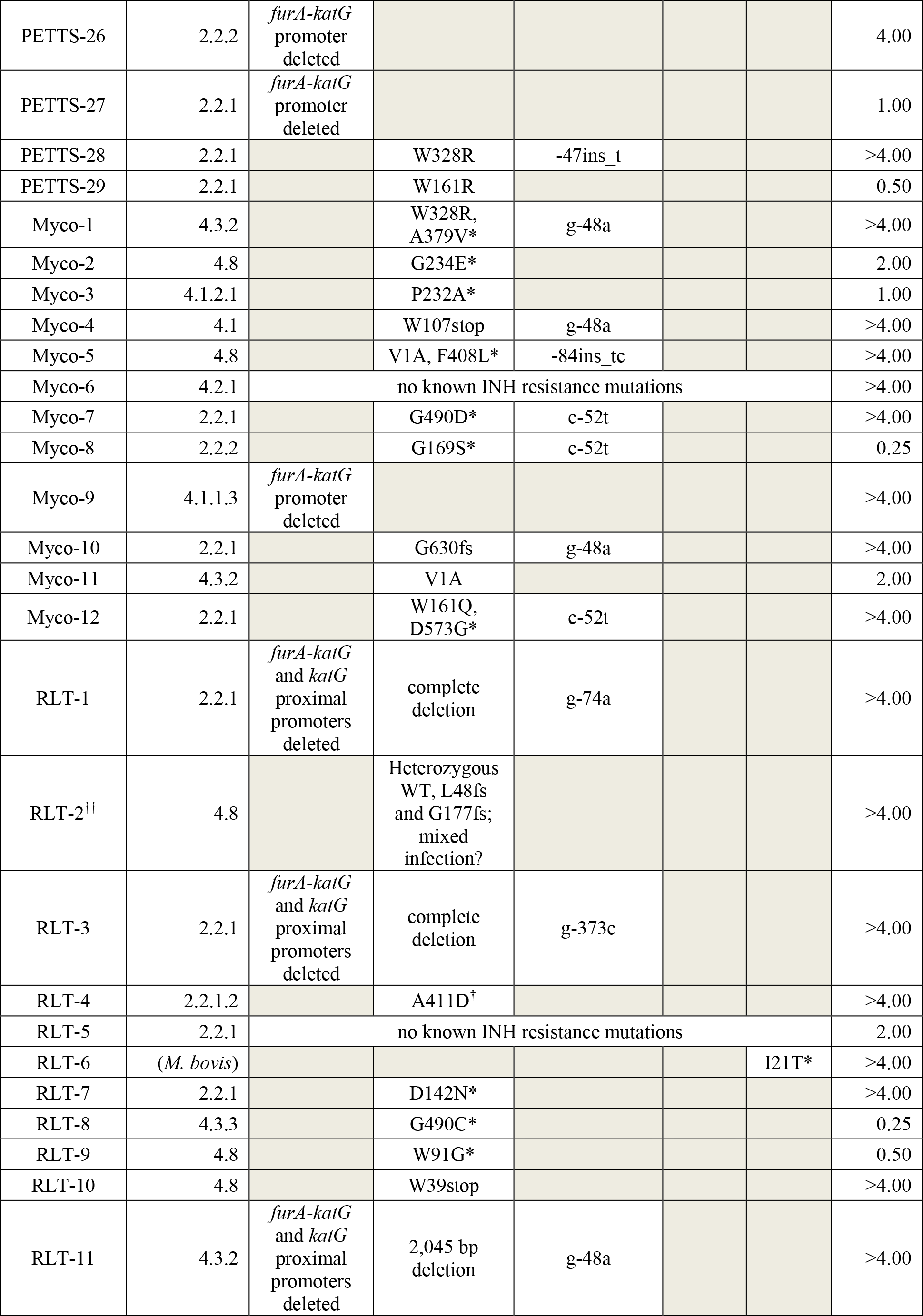
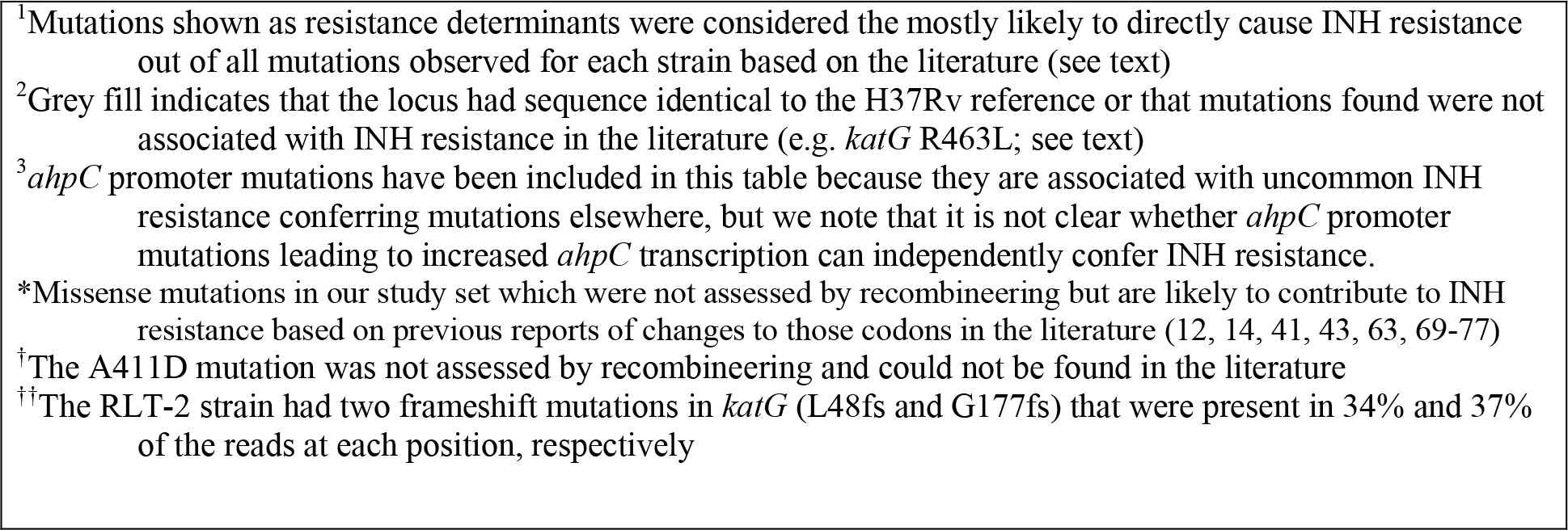
Uncommon mutations most likely^1^ to confer INH resistance in each clinical strain of the study set lacking canonical INH associated mutations, as identified by WGS

| Strain | *M. tuberculosis* SNP-based sublineage | INH resistance associated locus<sup>2</sup> |  |  |  |  | INH MIC (µg/mL) |
| --- | --- | --- | --- | --- | --- | --- | --- |
|  |  | *furA-katG* and proximal *katG* promoter regions | *katG* | *ahpC* promoter region<sup>3</sup> | *fabG1* | *inhA* |  |
| PETTS-1 | 1.2.2 |  |  | g-142a | L203L |  | 0.50 |
| PETTS-2 | 4.3.4.2 |  | 3' deletion | -47ins_t |  |  | >4.00 |
| PETTS-3 | 1.2.1 |  |  | g-142a | L203L | I21T* | 4.00 |
| PETTS-4 | 4.3.4.1 |  | V423fs | c-81t |  |  | >4.00 |
| PETTS-5 | 1.2.1 |  | S315N | g-142a |  |  | 1.00 |
| PETTS-6 | 1.2.1 |  | W300R | g-142a, g-74a |  |  | 0.50 |
| PETTS-7 | 1.2.1 |  | N138S | g-142a, g-48a |  |  | >4.00 |
| PETTS-8 | 1.2.1 |  | V1A | g-142a, g-51a |  |  | 1.00 |
| PETTS-9 | 2.2.1 |  | S315G |  |  |  | 0.50 |
| PETTS-10 | 1.2.1 |  | L415P | g-142a, c-54t |  |  | >4.00 |
| PETTS-11 | 1.2.2 |  |  | g-142a | L203L |  | 0.50 |
| PETTS-12 | 4.1.1.3 |  | E402stop | c-57t |  |  | >4.00 |
| PETTS-13 | 2.2.1 |  | V1A | c-52t |  |  | >4.00 |
| PETTS-14 | 2.2.1.1 |  | G285R* |  |  |  | 0.25 |
| PETTS-15 | 1.2.2 |  |  | g-142a | L203L | I194T* | 2.00 |
| PETTS-16 | 1.2.2 |  |  | g-142a | L203L | I194T* | 2.00 |
| PETTS-17 | 1.2.2 |  |  |  | L203L | I194T* | 4.00 |
| PETTS-18 | 1.2.2 |  |  | g-142a | L203L | I194T* | 2.00 |
| PETTS-19 | 1.2.2 |  |  |  | L203L | I194T* | 2.00 |
| PETTS-20 | 3.1.1 |  | S331fs | g-88a |  |  | >4.00 |
| PETTS-21 | 2.2.1 |  |  |  | L203L |  | 0.25 |
| PETTS-22 | 4.3.4.2.1 | no known INH resistance mutations |  |  |  |  | >4.00 |
| PETTS-23 | 2.2.1 |  | A480del | -47ins_t |  |  | >4.00 |
| PETTS-24 | 2.2.1 |  | G123fs | *ahpC-ahpD* overrepresented in sequence data |  |  | >4.00 |
| PETTS-25 | 2.2.2 |  | T322M* |  |  |  | >4.00 |
| PETTS-26 | 2.2.2 | *furA-katG*<br>promoter<br>deleted |  |  |  |  | 4.00 |
| PETTS-27 | 2.2.1 | *furA-katG*<br>promoter<br>deleted |  |  |  |  | 1.00 |
| PETTS-28 | 2.2.1 |  | W328R | -47ins_t |  |  | >4.00 |
| PETTS-29 | 2.2.1 |  | W161R |  |  |  | 0.50 |
| Myco-1 | 4.3.2 |  | W328R,<br>A379V* | g-48a |  |  | >4.00 |
| Myco-2 | 4.8 |  | G234E* |  |  |  | 2.00 |
| Myco-3 | 4.1.2.1 |  | P232A* |  |  |  | 1.00 |
| Myco-4 | 4.1 |  | W107stop | g-48a |  |  | >4.00 |
| Myco-5 | 4.8 |  | V1A, F408L* | -84ins_tc |  |  | >4.00 |
| Myco-6 | 4.2.1 | no known INH resistance mutations |  |  |  |  | >4.00 |
| Myco-7 | 2.2.1 |  | G490D* | c-52t |  |  | >4.00 |
| Myco-8 | 2.2.2 |  | G169S* | c-52t |  |  | 0.25 |
| Myco-9 | 4.1.1.3 | *furA-katG*<br>promoter<br>deleted |  |  |  |  | >4.00 |
| Myco-10 | 2.2.1 |  | G630fs | g-48a |  |  | >4.00 |
| Myco-11 | 4.3.2 |  | V1A |  |  |  | 2.00 |
| Myco-12 | 2.2.1 |  | W161Q,<br>D573G* | c-52t |  |  | >4.00 |
| RLT-1 | 2.2.1 | *furA-katG*<br>and *katG*<br>proximal<br>promoters<br>deleted | complete<br>deletion | g-74a |  |  | >4.00 |
| RLT-2<sup>††</sup> | 4.8 |  | Heterozygous<br>WT, L48fs<br>and G177fs;<br>mixed<br>infection? |  |  |  | >4.00 |
| RLT-3 | 2.2.1 | *furA-katG*<br>and *katG*<br>proximal<br>promoters<br>deleted | complete<br>deletion | g-373c |  |  | >4.00 |
| RLT-4 | 2.2.1.2 |  | A411D<sup>†</sup> |  |  |  | >4.00 |
| RLT-5 | 2.2.1 | no known INH resistance mutations |  |  |  |  | 2.00 |
| RLT-6 | (*M. bovis*) |  |  |  |  | I21T* | >4.00 |
| RLT-7 | 2.2.1 |  | D142N* |  |  |  | >4.00 |
| RLT-8 | 4.3.3 |  | G490C* |  |  |  | 0.25 |
| RLT-9 | 4.8 |  | W91G* |  |  |  | 0.50 |
| RLT-10 | 4.8 |  | W39stop |  |  |  | >4.00 |
| RLT-11 | 4.3.2 | *furA-katG*<br>and *katG*<br>proximal<br>promoters<br>deleted | 2,045 bp<br>deletion | g-48a |  |  | >4.00 |

<sup>1</sup>Mutations shown as resistance determinants were considered the mostly likely to directly cause INH resistance out of all mutations observed for each strain based on the literature (see text)
<sup>2</sup>Grey fill indicates that the locus had sequence identical to the H37Rv reference or that mutations found were not associated with INH resistance in the literature (e.g. *katG* R463L; see text)
<sup>3</sup>*ahpC* promoter mutations have been included in this table because they are associated with uncommon INH resistance conferring mutations elsewhere, but we note that it is not clear whether *ahpC* promoter mutations leading to increased *ahpC* transcription can independently confer INH resistance.
\*Missense mutations in our study set which were not assessed by recombineering but are likely to contribute to INH resistance based on previous reports of changes to those codons in the literature (12, 14, 41, 43, 63, 69-77)
<sup>†</sup>The A411D mutation was not assessed by recombineering and could not be found in the literature
<sup>††</sup>The RLT-2 strain had two frameshift mutations in *katG* (L48fs and G177fs) that were present in 34% and 37% of the reads at each position, respectively

### INH DST and MIC determination

All clinical strains were initially tested for INH susceptibility on Middlebrook 7H10 agar by the indirect agar proportion method as described by the Clinical and Laboratory Standards Institute (35). Resistance was defined as growth of >1% of the inoculum on a critical concentration of INH (0.2 μg/mL) as compared to a drug-free control. The MIC for INH was determined for all strains using the Sensititre MYCOTB MIC Plate (ThermoFisher Scientific) modified from a previous study (36). Briefly, Lowenstein-Jensen (LJ) slants were inoculated with 100 μL of frozen stock and incubated for no more than five weeks at 37°C. Growth was removed from the LJ slants with a pre-moistened sterile swab and resuspended in saline (0.8% w/v)/tween (0.2% v/v) solution with glass beads and briefly vortexed. The cell suspension was adjusted to a McFarland standard of ~0.5 and allowed to settle for 15 minutes. This suspension was diluted 1:100 in 10 mL of Middlebrook 7H9 with OADC (Thermo Scientific). After vortexing, the dilution tubes were allowed to settle for five minutes to minimize aerosols. Each 7H9 dilution was poured into a sterile reservoir and pipetted up and down three to four times to fully homogenize, then each well of the Sensititre plate was inoculated with 100 μL of the cell suspension. Plates were sealed with adhesive film, sealed in a plastic bag, and incubated at 37°C. Plates were read for MIC values at 10 and 21 days post-inoculation using the Sensititre Vizion Digital MIC Viewing System (ThermoFisher Scientific). The MIC was recorded as the lowest antibiotic concentration that reduced visible growth as compared to the drug-free control wells. Bitmap picture files of each plate reading were captured by SWIN software and read for a second time by the same operator on a separate computer to verify initial results. All reported MICs represent the mode from at least three independent experiments.

### WGS and analysis

MTBC strains were grown in Middlebrook 7H9 broth (6 mL) until culture turbidity was between OD_600_ 0.4 and 1.0. When possible, 200 μL of frozen stock were used instead of broth culture to minimize the chance of mutations occurring during *in vitro* growth. Genomic DNA was extracted using the ZR Fungal/Bacterial DNA MiniPrep Kit (Zymo), eluted in 25-30 μL of elution buffer, and quantified using a Qubit dsDNA BR or HS Assay Kit (ThermoFisher Scientific). Genomic DNA was sheared to a target size of 800 bp using an M220 Focused-Ultrasonicator and Holder XTU (Covaris), and libraries were prepared (25 ng input DNA per sample) using the Ovation Ultralow v2 Kit (NuGen) according to the manufacturer’s instructions. Libraries were pooled to 8 pM and sequenced on a MiSeq using the 500-Cycle MiSeq Reagent Kit v2 (Illumina). Sequenced genomes had an average median coverage of 118x and Q30 scores of 80%-90%. FASTQ file output from the MiSeq sequencing runs was assembled to the H37Rv reference strain (NCBI accession number NC_000962.3) using Lasergene SeqMan NGen (DNASTAR).

SNP calling and alignment parameters were set to default parameters in SeqMan NGen. Assemblies were visualized using Lasergene SeqMan Pro (DNASTAR). Single nucleotide polymorphisms (SNPs), small insertions/deletions (indels), and multi-kb indels were detected in SeqMan Pro using the SNP Report, Alignment View, Strategy View, Coverage Report, and Structural Variation Report tools. SNPs with a read depth <20 or a SNP % <25 were ignored. Trimmed sequences were aligned against H37Rv (https://www.ncbi.nlm.nih.gov/; accession number NC_000962.3) in BLAST (https://blast.ncbi.nlm.nih.gov/Blast.cgi) to determine the nature of insertions/deletions. Common INH-associated genes *katG (Rv1908c)*, *inhA (Rv1484)*, *furA (Rv1909c)*, *fabG1 (Rv1483)*, *ahpC (Rv2428)*, and other INH-associated loci found in the literature [*kasA (Rv2245)*, *srmR (Rv2242)*, *ndh (Rv1854c)*, *iniB (Rv0341)*, *iniA (Rv0342)*, *iniC (Rv0343)*, *Rv0340*, *nat (Rv3566c)*, *Rv1592c*, *fadE24 (Rv3139)*, *Rv1772*, *efpA (Rv2846c)*, *fabD (Rv2243)*, *accD6 (Rv2247)*, or *fbpC (Rv0129)*(37)] were searched for modifications. To determine the approximate level of gene duplication in the PETTS-24 strain at the *ahpC* region, the highest and lowest points of coverage depth were located in SeqMan Pro’s Strategy View and quantified in SeqMan Pro’s Alignment View.

### Preparation of electrocompetent, recombinant *M. tuberculosis*

H37Rv carrying the pJV128 recombineering plasmid was generated previously (34) and was used as a recipient strain in recombineering experiments as described therein, with some modifications. 7H9-succinate broth was prepared as described in (38) but cyclohexamide and carbenicillin were omitted. Aliquots (~120 μL) of electrocompetent cells were snap frozen in a dry ice/ethanol bath for three minutes and stored at −70°C until used.

### Recombineering oligo design and electroporation into H37Rv pJV128

All oligos used to generate mutations of interest were designed to anneal to the lagging strand of the H37Rv genome and were compared against H37Rv reference sequence (https://www.ncbi.nlm.nih.gov/; accession number NC_000962.3) to check for errors using BLAST (https://blast.ncbi.nlm.nih.gov/Blast.cgi). Oligos were ordered through Integrated DNA Technologies (IDT) in 100 nmole quantities with polyacrylamide gel electrophoresis (PAGE) purification. Experimental oligos were resuspended in molecular grade ddH_2_O to a concentration of 100 μM, and a working stock was made at 5 μM. The HygFix_For oligo, which is required for repair of the double stop codon in the pJV128 hygromycin (Hyg) B phosphotransferase gene, was prepared at a working stock concentration of 1 μM. One hundred ng of HygFix_For and 500 ng of experimental oligo (~10 μL total DNA volume) were premixed then added to 100 μL of room temperature electrocompetent cells. Reactions were electroporated on a Gene Pulser electroporator (Bio-Rad) under the following conditions: 2.5 kV, 1000Ω, 25 μF. After electroporation, all reactions were transferred to 1 mL of 7H9 broth and incubated shaking at 100 rpm for 48-72 hours to recover.

### Selection of plasmid-free recombinants for resistance-conferring mutations

Recovery cultures were serially diluted and spread plated onto 7H10 agar containing either Hyg (50 μg/mL) alone or Hyg (50 μg/mL) plus INH (0.2 μg/mL) to select for cells that were recombinant and had repaired the double stop codon mutation in *hygS*_amber_ on the pJV128 plasmid (Hyg only), or that had also gained an INH resistance-conferring mutation (Hyg+INH). Plates were incubated for three to four weeks before counting colonies. To allow for growth of cells that lost the pJV128 plasmid, four to ten colonies were picked from the 7H10 Hyg^50^+INH^0.2^ plates for each mutation and resuspended in 150 μL of plain 7H9 broth in a polystyrene 96-well plate. These plates were incubated for approximately one week, or until all wells were turbid. Thirty μL from each well were serially diluted in plain 7H9 broth and plated onto 7H10 agar containing 10% (v/v) sucrose. DNA was extracted from the remaining culture from each well and analyzed by Illumina WGS to check for i) the desired mutation and ii) the absence of confounding mutations and plasmid.

### Accession numbers

The MTBC clinical strain WGS data presented in this report is accessible at the Sequence Read Archive under accession number SRP137013 (https://www.ncbi.nlm.nih.gov/sra/SRP137013) and at BioProject under the accession number PRJNA448595 (http://www.ncbi.nlm.nih.gov/bioproject/448595).

## Results

### Selection of strains with discordant INH molecular and phenotypic results

We used clinical MTBC strains from previous studies stored in CDC’s Division of Tuberculosis Elimination Laboratory Branch to look for mutations that might explain discordant INH resistance test results. To build our study set, we evaluated 1278 international MDR-, pre-XDR-, and XDR-TB strains from the PETTS archive (39), 212 INH-R strains from a previous drug resistance study (31), and 12 INH-R strains from recent submissions to the MDDR service offered at CDC. All strains were isolated from clinical specimens between the years 2000 and 2017. To be eligible for study inclusion, strains were required to be phenotypically resistant to INH but lack the *katG* S315T and *fabG1-inhA* promoter mutations (c-15t, a-16g, t-8c, and t-8a) as determined by the Hain Genotype MTBDR*plus* v2 LPA test or Sanger/pyrosequencing (61/1502 strains). Strains were also excluded if unable to recover from frozen stock, if they tested susceptible to INH by Sensititre, or if a common INH resistance mutation (mentioned above) was discovered by WGS (Figure 1). Fifty-two strains met the criteria for inclusion. After SNP-based phylogenetic analysis, we determined that 51/52 strains were *M. tuberculosis* and one strain was *M. bovis.* Thirteen out of the 51 *Mtb* strains were of the Indo-Oceanic lineage (L1), 20/51 were of the East-Asian lineage (L2), 1/51 was of the East-African-Indian lineage (L3), and 17/51 were of the Euro-American lineage (L4) (Table 1).

### MIC determination for each strain

We determined the INH MIC for each strain using Trek Sensititre MYCOTB plates (Table 1 and **Supplemental Dataset S1**). Given that the Sensititre INH MIC of H37Rv is 0.6 μg/mL and two-fold differences in serial dilution MIC assays are unlikely to be significant, we considered strains with an MIC ≥0.25 μg/mL to be INH-R. The majority of the strains (39/52) were highly resistant to INH (MIC ≥2 μg/mL). Of these, 29/39 were resistant to the highest concentration tested (4 μg/mL). The remaining strains (13/52) expressed low-level INH resistance (MIC ≥0.25 but ≤1.00 μg/mL).

### WGS analysis to determine mutations linked to INH-R phenotype

In order to determine the genetic mechanisms leading to INH resistance in discordant strains of MTBC, we performed WGS on each of the clinical isolates. We reviewed available literature on INH resistance in MTBC (12-15, 18, 19, 22-24, 40-47) and works cited therein to deduce the most probable causes of this phenotype for each strain, which are listed in Table 1. Broadly, such mutations in our study set arepredicted to decrease KatG activation of INH prodrug or overexpress/modify the target of active INH, InhA.

In 26/52 strains, the INH-R phenotype could be explained by loss of *katG* due to known resistance-conferring mutations such as large deletions that removed *katG* promoter(s) and/or ORF sequence (n=7; Figure 2); frameshifts (n=5), nonsense mutations (n=3), or S315N,G missense mutations in *katG* (n=2); and the *fabG1* L203L mutation (n=9). Most (6/9) of the *fabG1* L203L mutations co-occurred with *inhA* ORF missense mutations. In contrast, 23/52 strains’ INH resistance was likely due to uncommon missense mutations in *katG* (n=21), a small indel in *katG* (n=1), or a missense mutation in *inhA* (n=1). Though these mutations are not yet considered high-confidence INH resistance markers, we were able to find supporting evidence for most of them as INH resistance determinants in the literature (see footnotes in Table 1) or validate them by functional genetics in *Mtb* (see below). The *katG* R463L mutation was also present in 33/52 strains, however this polymorphism is known to be a phylogenetic marker (32) and does not confer INH resistance (48). In 3/52 strains, no changes occurred that could readily explain the INH-R phenotype (Table 1 and **Supplemental Table S3**).

**Figure 2:**
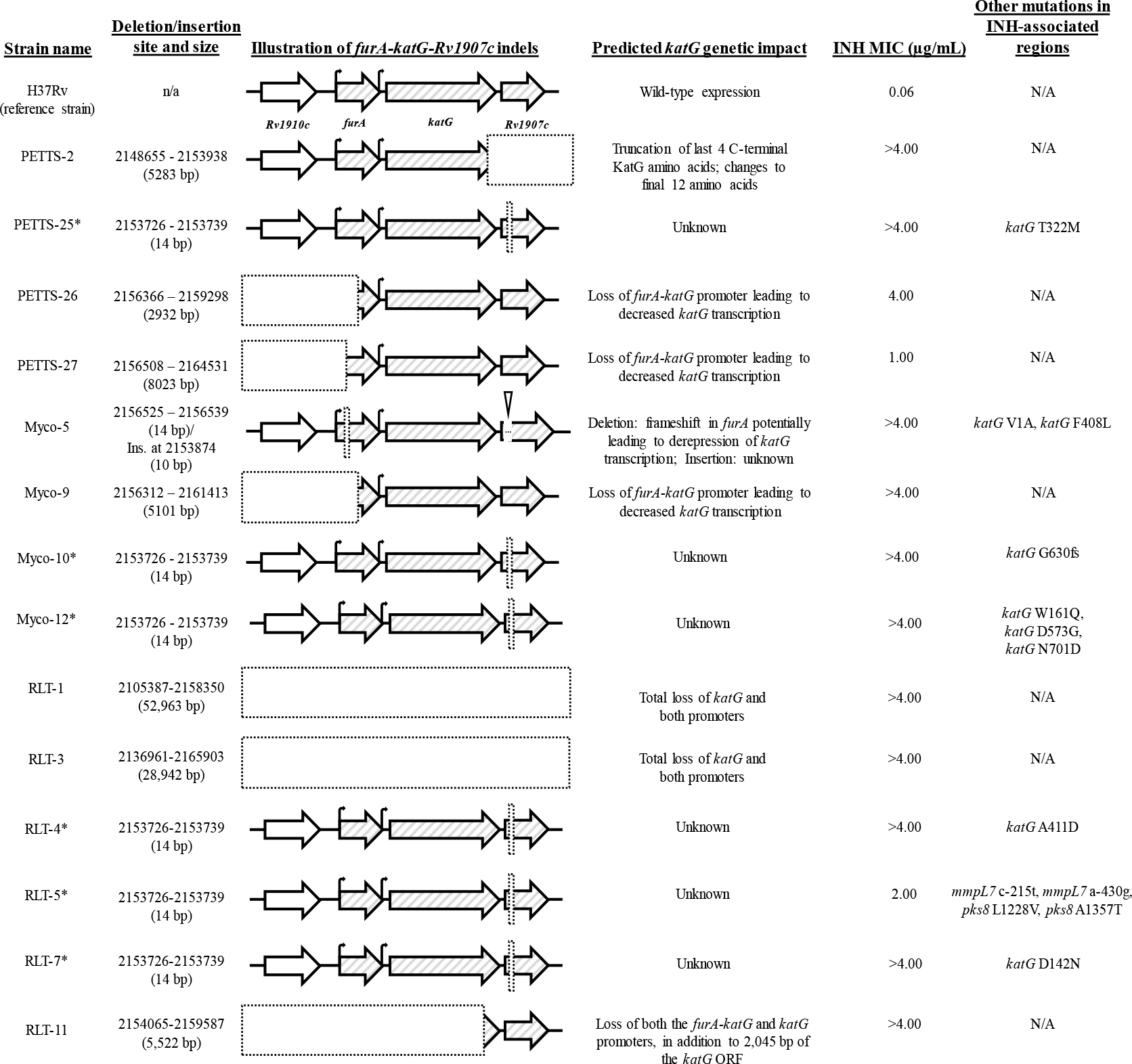
Genetic impact of large insertions and deletions at the *furA-katG-Rv1907c* operon in clinical MTBC strains. Of the 14/52 study set clinical strains that encoded large (>10 bp) insertions or deletions at the *furA-katG-Rv1907c* operon, 7/14 were predicted to impact *katG* activity leading to INH resistance. The insertions/deletions present at this locus in the other 7/14 strains were not predicted to impact INH resistance based on the literature, but were generally accompanied by mutations in *katG* (6/7 strains) likely to confer the INH-R phenotype. These seven strains also shared an identical 14 bp deletion in *Rv1907c* (marked with an asterisk; deleted sequence: TCATCCCCGTCTCG) and were of the same lineage (L2). One strain presented here, RLT-5, did not encode any mutations clearly related to INH resistance. Dotted line boxes indicate deleted sequence. Carrots indicate insertion sites. Hatched open reading frames are co-transcribed in *Mtb* (67). Promoters are indicated by bent arrows (45).

In addition to at least one probable INH resistance conferring mutation, 29/52 strains also carried *ahpC* promoter mutations. None of the clinical strains in our study set carried *ahpC* promoter mutations in isolation. One strain, PETTS-24, even appeared to encode a novel mechanism for *ahpC* overexpression: upon visual analysis of WGS data, PETTS-24 had an overabundance of reads from nucleotides 2717792-2727891 (10.1 kb), which includes the *oxyR’-ahpC-ahpD* region. On average, reads mapping to just the *ahpC* ORF were ~8-11x more abundant than the median genome coverage. We noted that reads at each end of the 10.1 kb overrepresented region had unaligned nucleotides which shared homology with genome sequence flanking the opposite end of the overrepresented region 10.1 kb away, suggesting a possible tandem array (49).

### Rare and unprecedented mutations in *katG* tested individually for their role in INH resistance

As mentioned above, 23/52 clinical strains encoded uncommon mutations in INH-associated loci which have not been directly shown (e.g., by functional genetic experiments) to confer INH resistance. To determine if previously unexamined *katG* mutations in our study set conferred INH resistance in a susceptible background, we attempted—by recombineering (50)—to generate 16 unique mutations in the *katG* open reading frame. Of these, 6/16 mutations have been previously reported in INH-R clinical isolates (*katG* V1A, N138S, W300R, S315G, S315N, W328R), 8/16 mutations are unprecedented in the literature (W161Q, W161R, E402stop, L415P, A480del, G601delins_GG, A606P, N701D), and 2/16 were controls to confirm the efficacy of the recombineering method for our purposes (W107stop, S315T).

The *katG* mutations V1A, N138S, W161Q, W161R, W300R, S315G, S315N, S315T, W328R, E402stop, L415P, and A480del were successfully generated in the H37Rv background. In contrast, we were unable to generate the *katG* mutations W107stop, G601delins_GG, A606P, and N701D. Instead, these desired mutations were either coupled with a second mutation within *katG* or did not occur at all (data not shown). During the recombineering process, we also unintentionally generated a *katG* A110V mutant, which we included in subsequent studies. Prior to MIC testing, we closely analyzed the SNP reports for recombineered strains to ensure that no confounding mutations were present that could impact INH resistance. Of the seven unintentional SNPs we observed in 8/13 recombineered strains, none of them occurred within 2,000 bp of any INH-associated locus. Six of these seven SNPs were either intergenic or encoded synonymous mutations. The single nonsynonymous SNP, encoding a *fadD12* Q471R mutation, was detected in only one of the *katG* S315T clones. Since the *katG* S315T is already thoroughly established as an INH resistance mutation and was included as positive control, we concluded that the SNP in *fadD12* would have no bearing on our experimental results. Thus, we considered all clones of our recombineered H37Rv *katG* mutants to be free from confounding mutations that might impact DST MICs.

INH MICs for each recombineered H37Rv *katG* mutant are shown in Table 2. Recombineered strain MICs for other anti-TB drugs included on the Sensititre MYCOTB plate did not differ significantly from those of the H37Rv parent strain (≤2-fold change) which suggests that the increased drug resistance conferred by recombineered mutations was specific to INH (data not shown). Not including the H37Rv S315T control, eight *katG* mutants had high-level INH resistance (MIC ≥2 μg/mL) and four *katG* mutants had low-level INH resistance (MIC ≥0.25 and ≤1.00 μg/mL). To our knowledge, five of the mutations listed in Table 2 (*katG* W161Q, W161R, E402stop, L415P, and A480del) have not been previously reported.

**TABLE 2:** Rare katG mutations that confer INH resistance as determined by recombineering.

| Strain | INH MIC<sup>3</sup><br>(µg/mL) | Clinical interpretation | INH MIC(s) in clinical strains with mutation<sup>4</sup><br>(µg/mL) |
| --- | --- | --- | --- |
| H37Rv | 0.06 | Susceptible | N/A |
| H37Rv *katG* V1A | >4.00 | High-level resistance | 1.00<sup>PETTS-8</sup>, >4.00<sup>PETTS-13</sup>, >4.00<sup>MYCO-5</sup>, 2.00<sup>MYCO-11</sup> |
| H37Rv *katG* A110V | 0.25 | Low-level resistance | N/A<sup>†</sup> |
| H37Rv *katG* N138S | >4.00 | High-level resistance | >4.00<sup>PETTS-7</sup> |
| H37Rv *katG* W161Q<sup>1</sup> | 0.25 | Low-level resistance | >4.00<sup>MYCO-12</sup> |
| H37Rv *katG* W161R<sup>1</sup> | 4.00 | High-level resistance | 0.50<sup>PETTS-29</sup> |
| H37Rv *katG* W300R | 1.00 | Low-level resistance | 0.50<sup>PETTS-6</sup> |
| H37Rv *katG* S315G | 0.25 | Low-level resistance | 0.50<sup>PETTS-9</sup> |
| H37Rv *katG* S315N | >4.00 | High-level resistance | 1.00<sup>PETTS-5</sup> |
| H37Rv *katG* S315T<sup>2</sup> | >4.00 | High-level resistance | N/A |
| H37Rv *katG* W328R | >4.00 | High-level resistance | >4.00<sup>PETTS-28</sup>, >4.00<sup>MYCO-1</sup> |
| H37Rv *katG* E402stop<sup>1</sup> | >4.00 | High-level resistance | >4.00<sup>PETTS-12</sup> |
| H37Rv *katG* L415P<sup>1</sup> | >4.00 | High-level resistance | >4.00<sup>PETTS-10</sup> |
| H37Rv *katG* A480del<sup>1</sup> | >4.00 | High-level resistance | >4.00<sup>PETTS-23</sup> |

<sup>1</sup>Novel mutations found in this study<sup>2</sup>Control mutation that is widely known to confer high-level INH resistance; we were not able to recombineer the other control mutation, *katG* W107stop (see text)<sup>3</sup>MICs of the recombineered *katG* mutants of H37Rv<sup>4</sup>MICs of the clinical strains encoding mutations chosen for functional genetics analysis are in grey. Corresponding clinical strain names from **Table 1** are in superscript; note that many of the clinical strains had multiple INH resistance associated mutations or other mutations (e.g. *katG* R463L) and thus MICs may not match those of recombineered strains harboring only a single *katG* mutation<sup>†</sup>The *katG* A110V mutation did not occur in our study set of clinical strains, but was previously associated with INH MICs of <1 µg/mL in 6/7 tested clinical isolates (64)

Generally, the degree of INH resistance conferred by each *katG* mutation was consistent between recombineered mutants and their corresponding clinical strains (Table 2), though we note that clinical strains often had additional INH associated mutations that prevent making a clear comparison. Strangely, the *katG* W161R and S315N mutations conferred high-level resistance in H37Rv but not in their respective clinical strains, PETTS-29 and PETTS-5. PETTS-5 additionally encodes a *katG* S140N mutation (51) and both of these clinical strains encode a *katG* R463L mutation, which is a known phylogenetic marker (32). It is not clear if these or other coincident benign mutations can ameliorate the effects of INH-R mutations in *katG*.

### Clinical strains that did not encode any predicted resistance mutations in *furA, katG, fabG1, inhA*, or *ahpC* or had mixed *katG* alleles

We explored the genome sequences of these “unexplained” strains further to determine if other, more obscure mutations related to INH resistance were present. Briefly, we found changes in 13 different genes among the three unexplained strains that might reduce INH susceptibility based on the literature (*accE5*, *efpA*, *mmpL7*, *msrA*, *mymA*, *pknH*, *pknK*, *pks8*, *nuoC*, *nuoG*, *nuoM*, *Rv1592c*, *sdh1A/Rv0248c*). Notable mutations include i) a large indel ~110 bp upstream from the operon encoding the EfpA MFS-type efflux pump, ii) SNPs upstream of the *mmpL7* gene, which could impact expression of the encoded INH efflux pump, and iii) a large deletion in *msrA*, whose encoded methionine sulfoxide reductase uses up NAD(P)H which could impact competition between NAD(P)H and INH-NAD adduct for InhA binding. **Supplemental Table S3** summarizes all mutations of interest for these three strains and our rationale for suggesting a connection to INH resistance. Since we did not test any of these mutations individually, these strains will require further investigation before the specific mechanism(s) conferring their INH resistance can be determined. A fourth strain, RLT-2, had a heterogeneous population of *katG* reads suggesting a mixed sampling of *Mtb* bacilli from a single patient. Two separate single bp deletions were observed in *katG* that would likely eliminate protein function via frameshift [37/108 (34%) reads at nt position 2155969 encoding a L48fs mutation and 44/120 (37%) reads at nt position 2155581 encoding a G177fs mutation].

## Discussion

In this study, we observed phenotypic INH resistance among MTBC clinical strains lacking the common mutations *katG* S315T and *fabG1-inhA* t-8a, t-8c, c-15t, a-16g. Early detection of less common INH resistance mutations could allow patients to be placed on appropriate therapy more rapidly and decrease the chances of treatment failure and acquired drug resistance. For our study set, the results demonstrate that INH resistance mutations not detected by conventional molecular methods could have been detected in 94% (49/52) of study set strains by expanding regions examined for *fabG1-inhA* and *katG* and accounting for large deletions that include *katG* and/or its promoters. An illustration summarizing INH resistance mutations confirmed previously in *Mtb*, and those described in this work, is presented in Figure 3.

**Figure 3:**
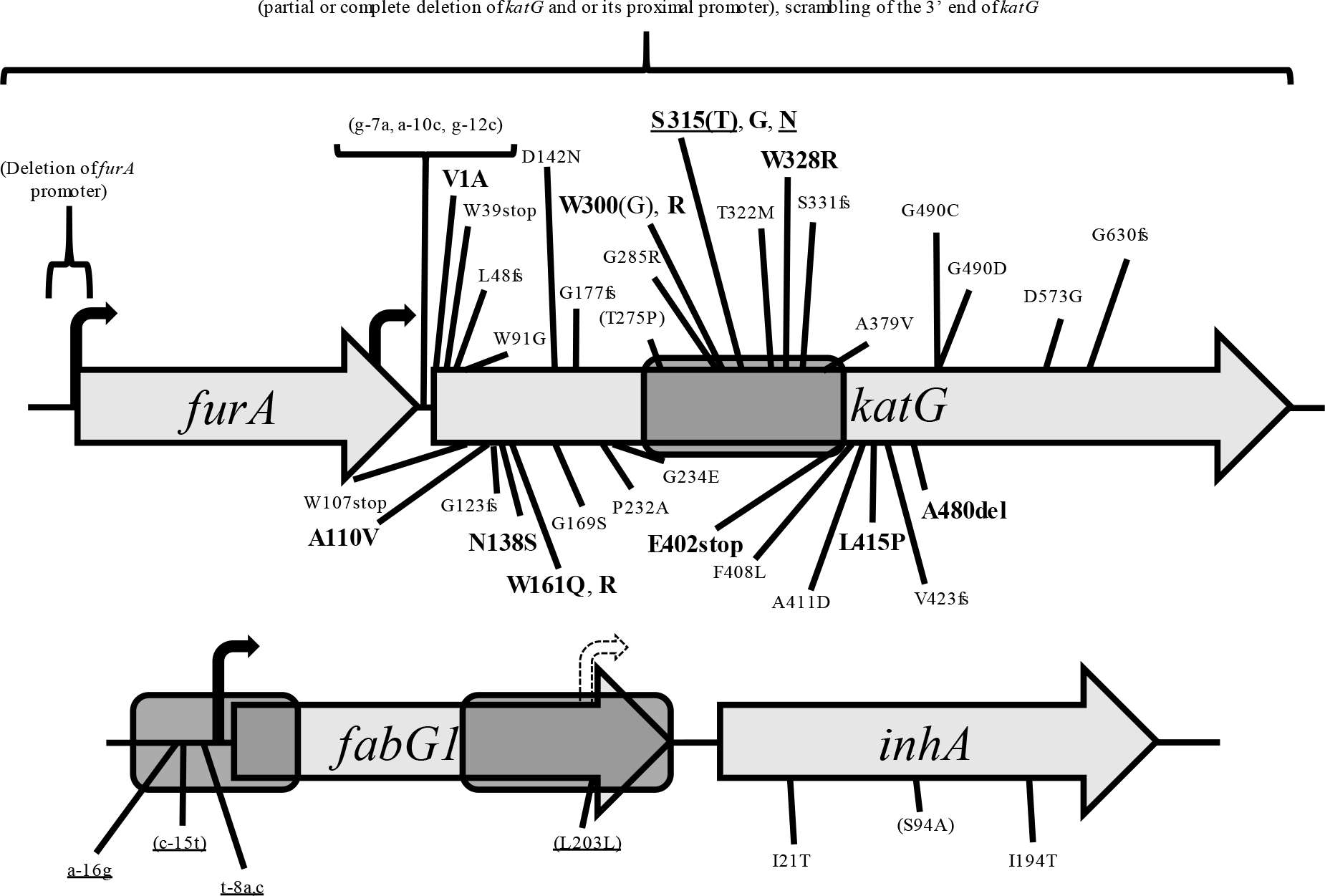
INH resistance determinants in *Mtb*. Illustration (not to scale) of the two primary loci involved in INH resistance, *furA-katG* and *fabG1-inhA*. Promoters are shown as bent arrows (a novel promoter generated by the *fabG1* L203L silent mutation is depicted with dashed outline). Shown in parentheses are mutations previously confirmed by functional genetics in *Mtb* (16–18, 21–23). Underlined mutations are targeted by conventional rapid molecular tests [Hain GenoType MTBDR*plus* v2 (29, 68), Nipro NTM+MDRTB Detection Kit 2 (30), CDC’s Molecular Detection of Drug Resistance (MDDR) service]. Windows of detection for CDC’s MDDR service Sanger sequencing assay, per current primer binding sites, are shown as grey bubbles (Jeff Driscoll, personal communication). Recombineered INH resistance mutations from this study are in larger text and bolded. Mutations in unformatted text were observed among INH-R clinical MTBC strains analyzed by WGS in this study and were generally considered likely to confer INH resistance due to previous reports in the literature (see Table 1). *ahpC* promoter mutations have not yet been definitively demonstrated to independently confer INH resistance in MTBC and thus are not depicted here. This illustration omits numerous mutations observed in clinical isolates of MTBC or evaluated in other organisms/biochemically, and is not intended to be comprehensive.

Although conventional molecular tests such as LPAs are rapid and specific, a recent meta-analysis found that the sensitivity of the Genotype MTBDR*plus* (Hain Lifescience) for the detection of INH resistance is highly variable depending on the study, and that pooled sensitivity across all included studies was only 83.4% (52). Even with the added sensitivity of targeted Sanger/pyrosequencing of the regions around *katG* S315 and the *inhA* promoter, it is likely that at least 9-10% of INH-R MTBC will be missed by these methods (31, 53). While small mutations such as SNPs or short indels could theoretically be detected by designing more comprehensive sequencing primers or adding mutation probes to future LPAs, these measures would require continual rounds of validation and would need periodic adjustment as new resistance determinants emerge. Larger mutations, like the multi-kb deletions we observed that are predicted to reduce or eliminate *katG* expression, would be even less practical to target with conventional methods due to the variability of their start and stop sites. Therefore, WGS is likely to be the foremost molecular DST method that can reliably detect all currently known and yet-to-be determined INH resistance mutations.

The *oxyR’-ahpC* intergenic region is another important locus associated with INH-R MTBC, though it is not targeted by either of the WHO-recommended LPAs. A recent meta-analysis of INH resistance mutations in clinical isolates found that among 24 publications released between the years 2000 and 2013 that reported sequencing the *ahpC* promoter region, the cumulative frequency of *ahpC* promoter mutations in INH-susceptible strains was 1.31%, but in INH-R strains was 8.88% (14). This lends support to the idea that some *ahpC* promoter mutations may be important markers of INH resistance (24, 54). It is interesting to note that 56% (29/52) of strains in our study set—which exclusively contains uncommon INH resistance markers—encoded changes to the *ahpC* promoter region. While we acknowledge that our study set was much smaller (52 strains) than that examining *ahpC* promoter mutations in the Seifert et al. study [~3000 strains; (14)], our results correlate *ahpC* promoter changes and INH resistance markers outside of *katG* S315T or the *inhA* promoter. Previous reports have also noted the association of *ahpC* promoter mutations with uncommon INH resistance genotypes (55, 56). We are currently working on generating some of the clinical *ahpC* promoter mutations from this study and hope the results of those experiments will bring clarity to the longstanding questions surrounding the importance of the AhpC alkylhydroperoxidase for evolution of INH resistance.

Notably, our WGS data for strain PETTS-24 revealed that SNPs and small indels are not the only mechanism by which *ahpC-ahpD* can be overexpressed. To our knowledge, this is the first reported instance of *ahpC* overexpression due to a predicted gene amplification event (57) resulting in a tandem array. However, it is possible that previous reports of *ahpC* overexpression in the absence of observable promoter mutations are due to a similar phenomenon. In *Mtb*, gene duplication events have been seen before (58–60), but to our knowledge are restricted to duplications and do not involve *ahpC*. No information regarding amplifications approaching 10x were found in the literature. Further investigation will be required to fully characterize this novel method of *ahpC* overexpression.

In total, our functional genetics experiments have confirmed the role of seven mutations in INH resistance and discovered five novel INH resistance mutations. However, we were unable to generate the *katG* G601delins_GG, A606P, and N701D mutations, suggesting that these mutations are unlikely to confer INH resistance. The failed recombinant clones frequently encoded a frameshift within 10 codons of the desired mutation in *katG* at positions that would fall within the recombineering oligo annealing site. We suspect that such frameshifts explain growth on INH selection during the recombineering process. Since the clinical strains encoding *katG* G601delins_GG (net result: an in-frame insertion of a codon encoding glycine), A606P, and N701D changes also had secondary mutations associated with INH resistance in the literature or that were shown to confer INH resistance in this work (*katG* W300R, W328R, D573G), we suspect G601delins_GG, A606P, and N701D are benign. However, there were no alternative mutations leading to INH resistance in the clinical strain with the *katG* W107stop mutation. It is not clear why the *katG* W107stop mutation could not be recombineered, though we note it has only been observed in the presence of *ahpC* promoter mutations [this work; (61)].

The recombineered *katG* mutations V1A, A110V, N138S, W161Q, W161R, W300R, S315G, S315N, S315T (positive control), W328R, E402stop, L415P, and A480del all independently conferred INH resistance in H37Rv. For the V1A and E402stop mutations, resistance was not surprising given that a GTG→GCG mutation would not be recognized as a start codon (62) and a nonsense mutation even at *katG* codon 454 was associated with complete loss of catalase activity (63). Likewise, N138S, W300R, S315G, S315N, W328R were also expected to increase the INH MIC since mutations at all of those residues are associated with decreased INH-NADH adduct (active drug) formation in biochemical studies of KatG enzyme (40). Notably, the W161Q, W161R, E402stop, L415P, and A480del mutations were not reported in the literature and A110V has only been reported in the presence of other confounding mutations (43, 64).

Since frameshift mutations occurred throughout *katG* in our INH-R failed recombinants, this observation taken together with our clinical and recombineered mutant data strongly supports the notion that targeted molecular DST methods should scan the entire *katG* ORF. The current model of exclusively using molecular diagnostics to target INH resistance mutations inside of “hotspots” may ultimately select for clinical strains that have non-canonical INH resistance mutations by depleting those with common ones from the host population. This scenario was already hypothesized by Torres and colleagues, who found >20 clinical *katG* mutations outside codon S315 by WGS of and used site-directed mutagenesis to show that several could independently confer INH resistance to *Mycobacterium smegmatis* (43). There is evidence to suggest that the rise of non-canonical INH resistance mutations is already happening in India, where there is a high burden of drug-resistant TB (65). WGS or expanded targeted sequencing could be used to detect such mutations.

Interestingly, 6% (3/52) of our study set strains expressed INH resistance that could not be explained by any known mechanisms. Upon WGS analysis, changes to genes implicated in INH efflux activity, cell wall composition, dormancy, redox control, and post-translational regulation of INH targets were observed (**Supplemental Table S3**). While we have yet to investigate any of these possible new INH resistance determinants by functional genetic experiments, our data suggest that future molecular tests may need to look outside of the known INH-associated genes (*furA*, *katG*, *fabG1*, *inhA*, *ahpC*, etc.) if all INH resistance is to be reliably detected. These results also demonstrate the utility of WGS for retrospective analysis of strains and validation of new resistance loci.

Knowledge of specifically low-level INH resistance mutations may allow for inclusion of high dose INH within a treatment regimen and limit cases where first-line therapy is not indicated. For example, the *katG* A110V, W161Q, W300R, and S315G mutations (MIC ≥0.25 but ≤1.00 μg/mL; Table 2), should (in the absence of other INH resistance determinants) allow for expanded treatment options. Indeed, the WHO updated their treatment guidelines for drug-resistant TB in 2016 to include high-dose INH for rifampin-resistant TB and MDR-TB in patients without suspected or confirmed high-level INH resistance (66). Thus, our finding that certain *katG* missense mutations by themselves confer only low-level INH resistance argues against a blanket policy removing INH from drug regimens in all cases of *katG* mutant TB.

## Conclusions

In summary, we have shown that clinical MTBC strains may express INH resistance in the absence of mutations targeted by conventional rapid molecular tests (LPAs, targeted sequencing). We demonstrated that INH resistance can be predicted in such strains using WGS. Several unproven and unreported mutations in *katG* were generated in the H37Rv genetic background and shown to independently confer low-or high-level INH resistance in *Mtb* using functional genetics. Our approach could be easily adapted to clarify resistance mechanisms for other drugs. Going forward, WGS or expansion of targeted sequencing will be crucial for rapidly detecting INH-R MTBC missed by conventional molecular methods.

## Acknowledgements

Use of trade names is for identification only and does not constitute endorsement by the US Department of Health and Human Services, the US Public Health Service, or the Centers for Disease Control and Prevention. The findings and conclusions in this report are those of the authors and do not necessarily represent the views of the Centers for Disease Control and Prevention. We thank the patients who gave their time and energy to contribute to this study, the clinical and microbiological staff at each of the enrollment sites in the PETTS study, and finally the public health professionals (CDC, Division of Tuberculosis Elimination, Laboratory Branch, Reference Laboratory Team) who archived and examined the clinical samples in the Mycobacteriology Laboratory Branch (MLB) and MDDR collections for their contributions to this work. We would also like to thank Melisa J. Willby and Glenn P. Morlock (CDC, Division of Tuberculosis Elimination, Laboratory Branch) for their excellent technical expertise with growth and extraction of DNA from MTBC bacteria, in addition to use of the Sensititre MYCOTB assay for DST, and Heather L. Alexander for her assistance with preliminary rapid molecular testing of clinical strains by line probe assay.

The Global PETTS Investigators include: Martie van der Walt, Jeannette Brand, South Africa Medical Research Council, Pretoria, South Africa; Thelma Tupasi, Janice Caoili, M. Tarcela Gler, Tropical Disease Foundation, Manila, Philippines; Carmen Contreras, Martin Yagui, Jaime Bayona, Socios en Salud Sucursal, Lima, Peru; Vaira Leimane, Liga Kuksa, Girts Skenders, State Infectology Centre of Latvia, TB and Lung Disease Clinic, Riga, Latvia; Laura E. Via, National Institutes of Allergy and Infectious Diseases, Bethesda, MD, USA; Soo Hee Hwang, National Masan Tuberculosis Hospital, Masan, Republic of Korea; Grigory V. Volchenkov, Tatiana Somova, Vladimir Oblast Tuberculosis Dispensary, Vladimir, Russian Federation; Somsak Akksilp, Wanpen Wattanaamornkiet, Wanlaya Sitti, Ministry of Public Health, Bangkok, Thailand; Hee Jin Kim, Chang-ki Kim, Korea Institute of Tuberculosis, Seoul, Republic of Korea; Boris Y. Kazennyy, Elena Kiryanova, Evgeniya Nemtsova, Orel Oblast Tuberculosis Dispensary, Orel, Russian Federation; Kai Kliiman, Tiina Kummik; Tartu University Lung Hospital, Tartu, Estonia; Piret Viiklepp, National TB Registry, Tallinn, Estonia; Ruwen Jou, Taiwan Centers for Disease Control and National TB Reference Laboratory, Taipei, Taiwan; Olga V. Demikhova, Larysa Chernousova, Central Tuberculosis Research Institute, Russian Academy of Medical Sciences, Moscow, Russian Federation; Ekaterina Kurbatova, Julia Ershova, Charlotte Kvasnovsky, Michael P. Chen, Melanie Wolfgang, U.S. Centers for Disease Control and Prevention, Atlanta, GA, USA.

## Funding Information

This project was supported in part by an appointment to the Research Participation Program at the Division of Tuberculosis Elimination (Laboratory Branch) in the National Center for HIV/AIDS, Viral Hepatitis, STD, and TB Prevention, Centers for Disease Control and Prevention, administered by the Oak Ridge Institute for Science and Education through an interagency agreement between the US Department of Energy and CDC. Matthew N. Ezewudo received support from a grant funded by the Bill & Melinda Gates Foundation, award number OPP1115887. PETTS clinical strains were derived from a study (39) funded by the US Agency for International Development, US Centers for Disease Control and Prevention, and US National Institute of Allergy and Infectious Diseases.

